# Molecular diversity and lineage commitment of human interneuron progenitors

**DOI:** 10.1101/2021.05.13.444045

**Authors:** Dmitry Velmeshev, Manideep Chavali, Tomasz J. Nowakowski, Mohini Bhade, Simone Mayer, Nitasha Goyal, Beatriz Alvarado, Walter Mancia, Shaohui Wang, Matthew Speir, Maximilian Haeussler, David Rowitch, Arturo Alvarez-Buylla, Eric J. Huang, Mercedes Paredes, Arnold Kriegstein

## Abstract

Cortical interneurons are indispensable for proper function of neocortical circuits. Changes in interneuron development and function are implicated in human disorders, such as autism spectrum disorder and epilepsy. In order to understand human-specific features of cortical development as well as the origins of neurodevelopmental disorders it is crucial to identify the molecular programs underlying human interneuron development and subtype specification. Recent studies have explored gene expression programs underlying mouse interneuron specification and maturation. We applied single-cell RNA sequencing to samples of second trimester human ganglionic eminence and developing cortex to identify molecularly defined subtypes of human interneuron progenitors and immature interneurons. In addition, we integrated this data from the developing human ganglionic eminences and neocortex with single-nucleus RNA-seq of adult cortical interneurons in order to elucidate dynamic molecular changes associated with commitment of progenitors and immature interneurons to mature interneuron subtypes. By comparing our data with published mouse single-cell genomic data, we discover a number of divergent gene expression programs that distinguish human interneuron progenitors from mouse. Moreover, we find that a number of transcription factors expressed during prenatal development become restricted to adult interneuron subtypes in the human but not the mouse, and these adult interneurons express species- and lineage-specific cell adhesion and synaptic genes. Therefore, our study highlights that despite the similarity of main principles of cortical interneuron development and lineage commitment between mouse and human, human interneuron genesis and subtype specification is guided by species-specific gene programs, contributing to human-specific features of cortical inhibitory interneurons.

## Main

The circuits in the mammalian brain contain two broad neuronal cell types: glutamatergic excitatory neurons and GABAergic inhibitory neurons called interneurons (IN). Interneurons are instrumental in controlling information flow in neuronal circuits and maintaining excitation-inhibition (E-I) balance (McBain and Fisahn, 2001). Disturbances of E-I balance and changes in specific IN subtypes have been associated with a number of neurodevelopmental and psychiatric disorders, including epilepsy (Sloviter, 1987), schizophrenia (Konradi et al., 2011) and autism spectrum disorder (Hashemi et al., 2017). Moreover, tipping the E-I balance toward excessive excitation has been shown to exacerbate neuronal cell death in neurodegenerative conditions and brain trauma by driving glutamate excitotoxicity (Zhang et al., 2016).

Unlike the progenitors of glutamatergic neurons that are located in the cortical ventricular and subventricular zones, neural stem cells generating the majority of cortical interneurons are located in the ganglionic eminences (GE) of the ventral telencephalon, with some interneurons produced in the preoptic area (Lavdas et al., 1999; Wichterle et al., 1999; Nery et al., 2002; Hansen et al., 2013; Ma et al., 2013). The GE is subdivided into three major regions: the medial (MGE), lateral (LGE) and caudal (CGE) ganglionic eminences. The progenitor cells of the MGE and CGE produce specific IN lineages that differentiate into defined cortical adult IN subtypes (Xu et al., 2004; Butt et al., 2005; Cobos et al., 2006). Recent single-cell RNA sequencing (scRNA-seq) studies of the GE and cortical interneurons in the mouse (Mayer et al., 2018; Mi et al., 2018) have revealed a model of IN lineage development whereby GE progenitors express lineagespecific transcription factors and produce committed immature INs that differentiate into adult IN subtypes upon migration to the cortex. However, it is currently not known to what degree IN progenitor and lineage-specific IN developmental programs identified in the mouse are reflective of the developing human brain.

To study the processes of human cortical IN production, lineage commitment and maturation, we collected scRNA-seq data from microdissected MGE, LGE and CGE tissue from six individuals during the second trimester spanning ages 11-20 postconceptional weeks (PCW) (**Figure 1A**), which coincides with the peak of IN production in the human brain (Hansen et al., 2013; Ma et al., 2013). In addition, we collected scRNA-seq data from interneurons migrating through the germinal and marginal zones and combined these data with previously published scRNA-seq data from cells in the human MGE and INs in the developing cortex (Nowakowski et al., 2017) (**Methods**). After filtering cell multiplets and low-quality cells, the final dataset included 2,241 single-cell transcriptomic profiles (**Table S1**). Unbiased clustering using Seurat (Butler et al., 2018) revealed 10 cell clusters (**Figure 1B**) and two major groups of cell types. The first group contained two clusters of IN progenitors that were classified as radial glia cells (RGC) and intermediate precursor cells (IPC) based on expression of known markers, such as vimentin (VIM) and nestin (NES) for RGCs and ASCL1 for IPCs (**Figure 1C**). We observed more IPCs in the MGE compared to the CGE, and the LGE contained significantly fewer RGCs and IPCs compared to both the MGE and the CGE (**Figure 1D**). The second major group of clusters contained postmitotic INs that expressed GAD1 and markers of neuroblast migration such as DCX and ARX. Remarkably, postmitotic interneurons that had not yet migrated to the cortex already clustered into three lineages: MGE-derived cortical INs (IN-MGE) expressing MAF and LHX6; CGE-derived cortical INs (IN-CGE) expressing NR2F2 (COUP-TFII) and PROX1; and striatal GABAeric neurons (GABA-STR) expressing MEIS2 (**Table S2**). We did not observe a similar segregation of RG and IPCs based on lineage, suggesting that the progenitor cells in the MGE and CGE are not as molecularly distinct as their neuronal progeny. The MGE and CGE-derived INs formed several transcriptomic clusters, with the clusters primarily containing cells from the cortex expressing the highest levels of lineage-specific transcription factors and pan-IN markers (**Figure 1D-E**). We observed two clusters of postmitotic interneurons that represented a transitional state between progenitors and lineage-committed INs. We reasoned that these clusters (nIN-1 and nIN-2), that did not express lineage-specific markers but stated to express interneuron markers, such as *GAD1*, contain newborn interneurons. Therefore, similar to the mouse, IN lineage commitment takes place in the ganglionic eminence in the developing human brain, and the MGE and CGE-derived INs become gradually more molecularly distinct as they migrate to the cortex.

**Figure 1.**
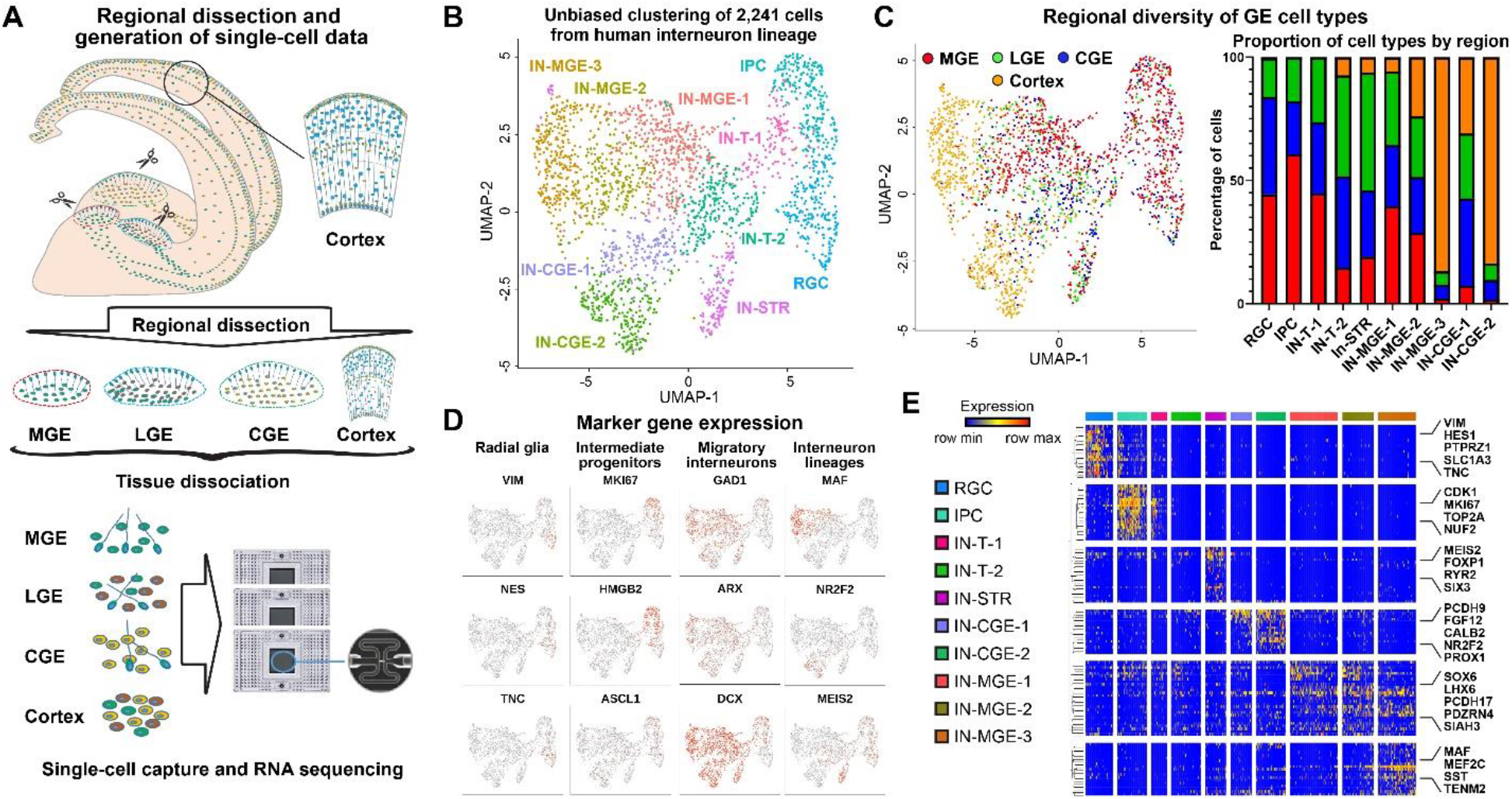
Single-cell RNA-seq analysis of human cortical interneurons during the second trimester of development. **A)** Cartoon depicting the experimental design and the brain regions sampled. **B)** Unbiased clustering of the scRNA-seq profiles obtained from human GE cells and cortical interneurons. RGC - radial glial cells, IPC - intermediate precursor cells, IN-T – newborn interneurons, IN-MGE – MGE-derived cortical interneurons, IN-CGE – CGE-derived cortical interneurons, IN-STR – striatal interneurons. **C)** Distribution of cells from the subregions of the GE and the cortex across transcriptomic clusters. **D)** Expression of markers of radial glia, intermediate progenitors, immature neurons and interneuron lineage-specific markers. **E)** Heatmap illustrating expression of top cluster enriched genes across all populations of cortical interneuron lineages during the second trimester.

We next asked whether distinct populations of IN progenitors are characterized by molecular signatures that enable them to produce lineage-committed INs. We performed differential expression analysis by comparing RG cells and IPCs from the MGE and CGE as well as from the cortex (**Figure 2A, Table S3**). We identified a number of genes enriched in MGE RG compared to CGE RG, and vice versa (**Figure 2B**). Among these genes, we observed canonical MGE and CGE transcription factors (TFs) NKX2-1 and NR2F2, as well as TFs that have not previously been reported to be enriched in MGE and CGE progenitors, such as TOX3, LMO1 and HEY1. Immunohistochemical (IHC) analysis of TOX3 protein expression in the human MGE revealed a cluster of NKX2-1-positive ventricular zone (VZ) cells at the MGE/LGE border that express TOX3 (**Figure 2C**), while it was not expressed by CGE VZ cells (**Figure 2D**). This suggests that some TFs are expressed in regionally restricted populations of ganglionic eminence radial glia. We also found a smaller number of region-specific neuronal? genes in IPCs, suggesting that lineage-specific gene programs are transiently repressed in IPCs and newborn INs before being upregulated in lineage-committed INs (**Figure 2F**). By performing Gene Ontology (GO) enrichment analysis of region-specific progenitor markers we found that these genes are enriched in processes associated with cell differentiation, regionalization, and transcriptional regulation (**Figure S1A**). This suggests that MGE and CGE progenitors are express region-restricted transcriptional regulators, which may prime production of specific IN lineages.

**Figure 2.**
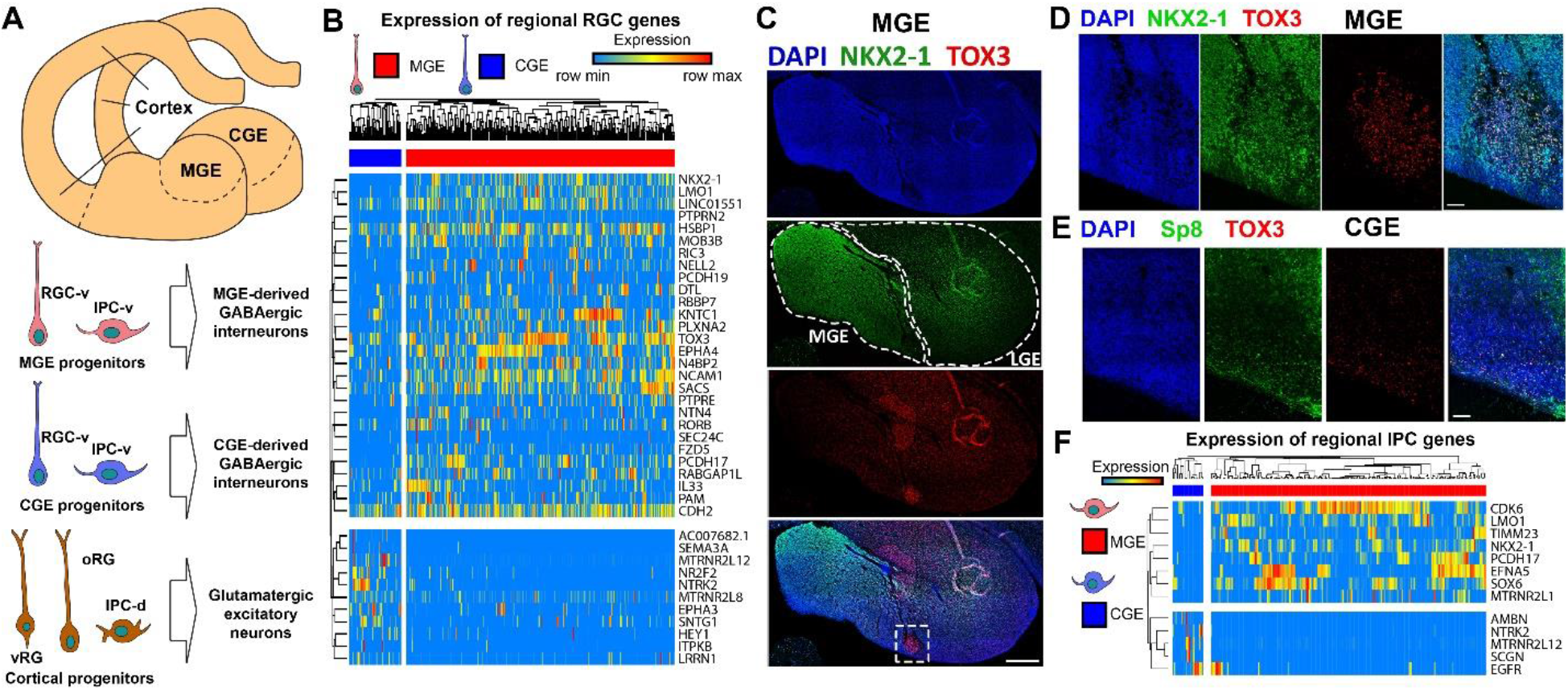
Region-specific signatures of GE progenitors. **A)** Overview of progenitor populations in the GE and the cortex as well as the neuronal lineages they produce. **B)** Heatmap of genes enriched in RGC populations in the MGE and CGE. **C)** Immunohistochemical staining for TOX in the human MGE and LGE. Scale bar: 500um. **D-E)** IHC for TOX3 in the human MGE and CGE. Scale bar: 50um. **F)** Genes enriched in MGE- and CGE IPCs.

In order to compare human RG cells and IPCs of the GE to dorsal cortical progenitors, we co-clustered dorsal and ventral progenitor cells (**Figure S1B**). This analysis further subdivided GE RG cells into two subpopulations that we named RGC-v-1 and RGC-v-2. We then identified gene markers of each ventral and dorsal progenitor cell type. By comparing these markers, we observed that ventral RG cells share more gene expression signatures with cortical ventricular radial glia (vRG) compared to outer radial glia (oRG) (Pollen et al., 2015) (**Figure S1C**). In addition to canonical ventral-specific DLX genes, ventral progenitors expressed novel markers, such as FLRT2 (expressed in ventral RG and IPCs) and LIX1 (expressed in ventral RG) (**Figure S1D-E**). DLX1 and DLX2 were expressed in both RG-v-2 and IPCs but not RG-v-1, while DLX5 and DLX6 were expressed specifically in IPCs. This suggested to us that RG-v-2 represents a transitional cell state between *bona fide* ventral RG and IPCs, and that a code of DLX genes regulates ventral progenitor differentiation. Therefore, regionalization and differentiation of ventral progenitors appear to be regulated by a coordinated expression of regional markers, including transcriptional regulators.

To test if transcriptional regulators specifically expressed in the human MGE or CGE exhibited regional specificity in the mouse, we performed clustering of scRNA-seq data from E13.5-E14.5 mouse GE and E18.5 mouse cortical interneurons (Mayer et al., 2018) (**Figure S2A-C**). We identified RG and IPC clusters, as well as lineage-committed postmitotic INs. We observed that a number of human progenitor markers were expressed in a region-specific manner in human but not mouse GE (**Figure S2D**), suggesting evolutionary emergence of novel mechanisms of GE progenitor regionalization.

In order to explore the molecular events underlying lineage commitment of human cortical INs, we performed trajectory analysis of the combined GE and cortical IN scRNA-seq data using Monocle (Cao et al., 2019) (**Figure 3A**). We observed that the trajectory graph first passed through RG, IPCs, and the cluster of newborn INs, after which it split into three branches. The branches corresponded to the MGE and CGE cortical IN lineages and the striatal GABAeric neuron lineage. We next calculated pseudotime values for each cell in the dataset by assigning the starting point of the trajectory to the terminal node in the RGC cluster (**Figure 3B**). By identifying genes that change their expression with pseudotime in either the MGE or CGE IN lineages, we discovered genes that are dynamically expressed during human cortical IN lineage commitment (**Figure 3C-D, Table S4**). These included known MGE and CGE-specific markers, as well as genes that have not been previously associated with IN commitment, such as the MGE-specific postsynaptic density protein AKAP5 and the CGE-specific Protein kinase C alpha (PRKCA). Hierarchical clustering revealed five modules of genes dynamically expressed in the MGE and CGE lineages (**Figure 3E**). Genes in these modules were expressed at different stages of IN differentiation and were associated with specific cellular processes. For example, the magenta module included genes expressed at low level in IPCs and newborn interneurons, but upregulated in lineage-committed INs (**Figure 3F**). These genes were associated with synaptic signaling, neurite outgrowth, adhesion and cell migration, and included known and novel lineage-specific genes, such as the axon guidance cue, SEMA6A, and the chemokine receptor, ACKR3, regulating cell migration. In contrast, genes in the red module were highly expressed in IPCs and to some degree in early newborn INs and were associated with cell division. These genes followed similar trajectories in both the MGE and CGE (**Figure 3G**). These findings strengthen the notion that cortical INs pass through a common undifferentiated state before committing to a specific lineage. To test whether some of the human IN lineage-specific dynamically expressed genes we identified were also expressed in the MGE and CGE lineages in the mouse, we performed trajectory and pseudotime analysis of mouse embryonic GE cells and cortical INs (**Figure S2E**). The canonical MGE and CGE-specific markers were expressed in a similar fashion in both mouse and human (**Figure S2F**). However, we identified a number of genes expressed during MGE and CGE lineage commitment in the human IN but not in the mouse (**Figure S2F, Table S4**), suggesting possible evolutionary changes in cortical IN differentiation programs. Additionally, in order to investigate whether MGE and CGE lineage INs migrate through different cortical laminae, we compared cortical interneurons captured from either the germinal (GZ) or marginal (MZ) zones in the developing cortex. We noticed that MGE INs were found in both the cortical GZ and MZ in the UMAP plot, whereas CGE INs separated into distinct groups migrating through the GZ or MZ (**Figure S3A**). By comparing the transcriptional profiles of CGE interneurons in the MZ and GZ, we identified a number of genes that differentiate these populations, including CNR1 and FGF12 in the MZ, and POGK and SKIL in the GZ (**Figure S3B**). These genes were either not expressed in the MGE lineage or not specific to the MZ or GZ. Our findings suggest that molecularly distinct populations of INs migrate through different cortical laminae. Finally, to explore dynamic gene expression during human and mouse striatal GABAergic lineage commitment, we applied Monocle to the cells in this lineage (**Figure S3C**). Contrary to cortical INs, we did not observe clear examples of dynamically expressed genes in this lineage that are specific to the human but not mouse (**Figure S3D**), suggesting differences in the evolution of gene programs for cortical and subcortical lineages.

**Figure 3.**
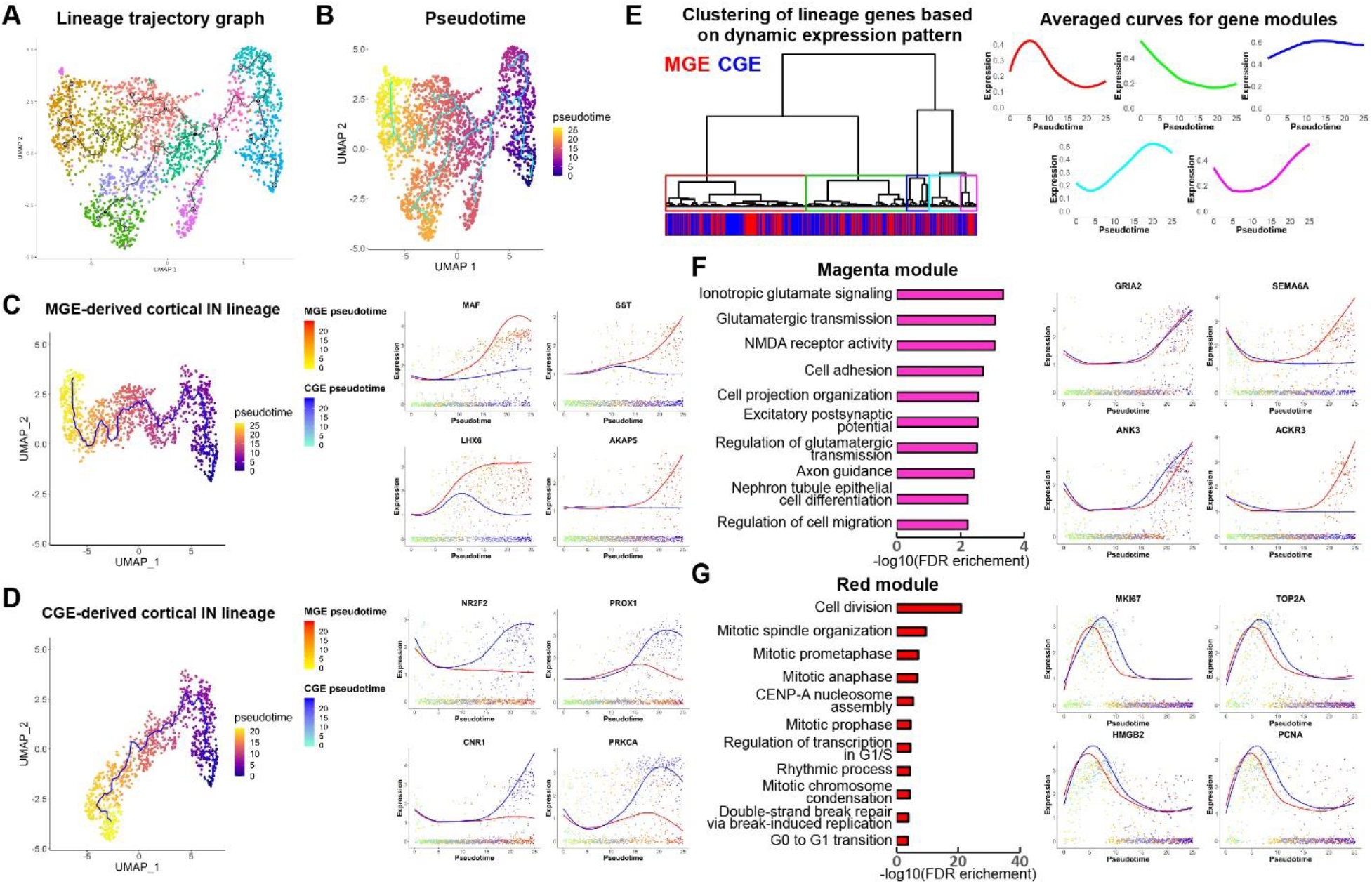
Lineage trajectory analysis of human interneuron lineages during the second trimester of development. **A)** Lineage trajectory graph for IN lineages. **B)** Pseudotime assignment of single cells in the lineages. **C)** MGE cortical interneuron lineage and lineage-specific genes expressed dynamically during lineage progression. **D)** CGE cortical IN lineage and lineage-specific pseudotime-dependent genes. **E)** Clustering of lineage-specific pseudotime-dependent genes according to their dynamic expression patterns as well as averaged expression curves for resulting gene modules. **F)** Gene ontology analysis of genes in the magenta module as well as shared and lineage-specific genes belonging to this module. **G)** Biological processes enriched for genes in the red module and curves of dynamic expression of red module genes in the MGE and CGE cortical interneuron lineages.

Developmental IN lineages give rise to a number of mature IN subtypes in the adult neocortex. In order to explore the molecular events underlying maturation of these subtypes, we integrated our developmental IN dataset with single-nucleus RNA-seq data from the adult human cortex of neurotypical individuals (Velmeshev et al., 2019) (**Figure 4A-B**). We identified clusters corresponding to GE progenitors, MGE and CGE-committed fetal interneurons, as well as four subtypes of adult cortical INs expressing somatotstatin (SST), parvalbumin (PV), vasoactive intestinal peptide (VIP) and the synaptic marker, SV2C, that has been shown to label neurogliaform cells (**Figure 4C**). We then performed trajectory and pseudotime analysis using the integrated dataset (**Figure 4D**) and focused on IN subtypes derived from MGE (SST and PV) and CGE (VIP and SV2C) lineages (**Figure S4A**). Adult INs expressed certain fetal lineage markers, such as LHX6 for SST and PV subtypes, and PROX1 for VIP and SV2C subtypes (**Figure 4E, Figure S4A**), confirming that these adult IN types originate from specific GE niches. By focusing on genes that are enriched in a specific cell subtype within the MGE and CGE lineages, we identified subtype-specific developmentally-regulated genes. These included transcription factors BACH1 and SOX5 for SST and PV INs (**Figure 4E**), as well as a VIP-specific noncoding RNA, PWRN1, located in the region of chromosome 15, that is imprinted in the Prader-Willi syndrome, and a SV2C-specific transcription factor, TOX (**Figure S4A**). Interestingly, a number of subtype-specific genes were initially expressed in progenitors and uncommitted INs, but later become restricted to a single IN subtype, suggesting that adult IN lineage commitment occurs early in development. We tested for expression of one such marker, BACH1, by performing IHC and single-molecule RNA FISH in human adolescent cortical tissue sections (**Figure 4F**), which validated enrichment of BACH1 in SST compared to PV interneurons. To test whether some adult human IN subtype-specific markers are also expressed in mouse INs, we performed an analogous analysis in the mouse by integrating fetal and adult IN scRNA-seq data followed by trajectory analysis (Yao et al., 2020). We identified a number of genes expressed in specific adult IN subtypes in the human but not in the mouse, including the WNT signaling inhibitor, DKK2, expressed in human SST INs, the transcriptional regulator, LDB2, expressed in PV INs (**Figure S3B**), as well as SEZ6L, implicated in epileptic seizures, and the transcriptional coactivator EYA4, expressed in VIP and SV2C interneurons (**Figure S3C**), respectively. Therefore, transcriptional regulation and modulation of signaling pathways during development can contribute to differences between human and mouse adult interneuron subtypes.

**Figure 4.**
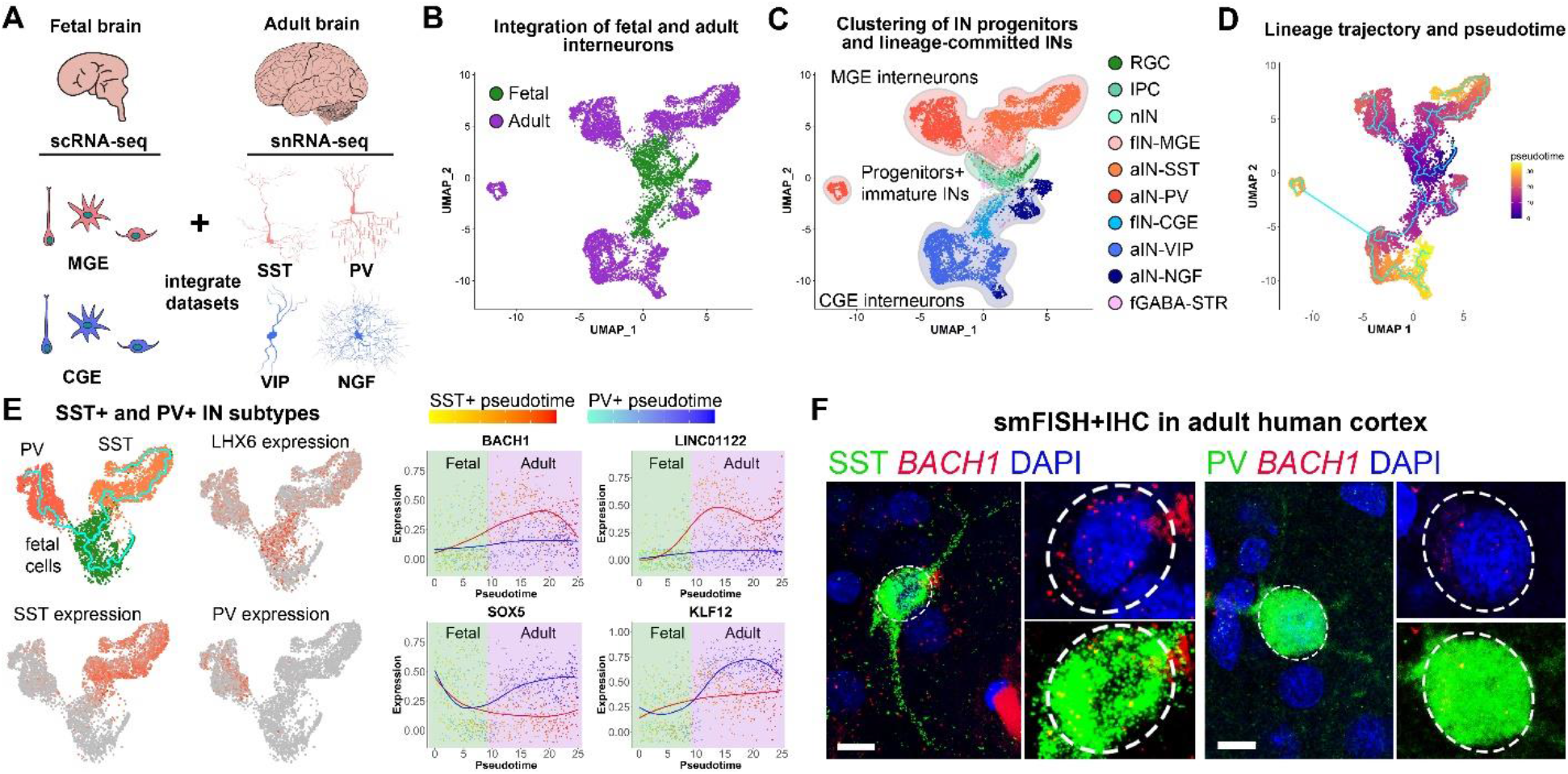
Early commitment of adult human cortical interneuron lineages. **A)** Cartoon depicting the major adult interneuron populations analyzed and the progenitor populations that produce them. **B)** Integration of fetal and adult interneuron datasets. **C)** Transcriptomic clusters corresponding to fetal and adult IN lineages and subtypes as well as to GE progenitors. nIN – newborn interneurons, fIN – fetal interneurons, aIN – adult interneurons, SST – somatostatin interneurons, PV – parvalbumin interneurons, VIP – VIP interneurons, NGF – neurogliaform cells. **D)** Lineage trajectory and pseudotime for the integrated IN datasets. **E)** Analysis of MGE-derived SST and PV interneuron commitment. SST- and PV-specific genes that are expressed during fetal development are highlighted on the right. **F)** Single-molecule RNA FISH for *BACH1* combined with IHC for SST and PV in postnatal human adult cortex. Scale bar: 10 um.

We performed a comprehensive single-cell analysis of human cortical interneuron development and compared it to the analogous process in the mouse. The human ganglionic eminence contains two main subtypes of progenitor cells: radial glia and intermediate progenitors. Interneuron lineage commitment can be observed in the GE before postmitotic interneurons migrate to the neocortex. Unlike postmitotic interneurons, GE progenitors do not segregate into lineage-committed populations based on their global gene expression profiles. However, the progenitor cells in the MGE and CGE express a small but highly restricted set of genes, including transcriptional regulators, that are region-specific in the human but not mouse. Some of these regulators, such as transcription factor TOX3, are restricted to a small population of GE progenitors, suggesting granular specification of human interneuron precursors. Newborn interneurons lose expression of region-specific markers before they start expressing lineage-specific markers characteristic of post-mitotic interneurons. Therefore, similar to the mouse brain, our findings suggest that a small number of transcriptional and epigenetic regulators may prime regional human interneuron progenitors toward producing interneurons of specific lineages. These regulators may in turn activate specific differentiation programs in committed interneurons as they mature. We find that despite the main principles governing interneuron lineage commitment in the mammalian neocortex being conserved between the mouse and human, a number of genes, including transcriptional regulators, demonstrate regional and lineage-specific expression in human ganglionic eminence progenitors but not in the mouse. These novel regulators may contribute to increased diversity of interneuron progenitors in the human ganglionic eminence by defining subpopulations of progenitors not present in the mouse. Additionally, we identify gene programs underlying gradual commitment of interneurons to specific developmental lineages and adult subtypes. Among these, we find a number of cell migration, cell adhesion, and synaptic genes that are upregulated during the development of specific human but not mouse interneuron lineages. We identify genes expressed in interneurons in the developing human but not mouse brain that become restricted to adult interneuron subtypes, such as transcription factor BACH1 in human somatostatin interneurons. This finding suggests that the process of adult human interneuron subtype specification utilizes regulatory mechanisms that are different from the mouse, potentially contributing to human-specific molecular and functional features of human GABAergic interneurons. This finding suggests that evolutionary changes in interneuron development may be associated both with the specification of progenitor domains and the integration of interneurons into cortical circuits. Previous comparative studies of mature cell types in human and mouse cortex suggested a general conservation of cell types between the two species while observing species-specific molecular programs defining these cell types (Hodge et al., 2019). We find a similar phenomenon during interneuron development, where the main lineages and developmental principles are conserved between the human and the mouse, but the gene regulatory programs diverge to accommodate species-specific requirements for neuronal function. We believe that our study provides novel findings that will contribute to understanding human brain development and neurodevelopmental disorders, as well as to creating better human cellular models of neural development by benchmarking human-specific developmental gene programs.

## Acknowledgements

We thank Dr. Cory Nicolas for discussion of the project and comments on the manuscript.

We thank the NIH for funding support of this project, in particular for grants R35NS097305, U01MH114825 and U01MH114825 awarded to Arnold Kriegstein and K99MH121534 awarded to Dmitry Velmeshev.

## Author Contributions

D.V. designed the study, collected single-cell RNA-seq data, performed data analysis and wrote the paper. M.C. performed RNAscope experiments. T.N. performed dissection of the MGE, CGE and LGE. M.B. performed IHC experiments. S.M. performed RNAscope experiments. N.G. performed imaging of single cells captured on C1 chips. B.A. helped with single-cell RNA-seq library preparation. W.M. helped with IHC experiments. S.W. collected developing human tissue samples. M.S. and M.H. designed the web browser and deposited the data. D.R. and A.A-B helped with manuscript preparation. E.H. and M.P. provided adolescent human brain tissue samples and helped with manuscript preparation. A.K. designed and managed the project and prepared the manuscript. All authors read the manuscript.

## Competing interests

A.A.-B. is the Heather and Melanie Muss Endowed Chair in Neurological Surgery at UCSF and is Co-founder and on the Scientific Advisory Board of Neurona Therapeutics. A.K. is Co-founder, consultant and on the Scientific Advisory Board of Neurona Therapeutics.

---

**Figure S1.**
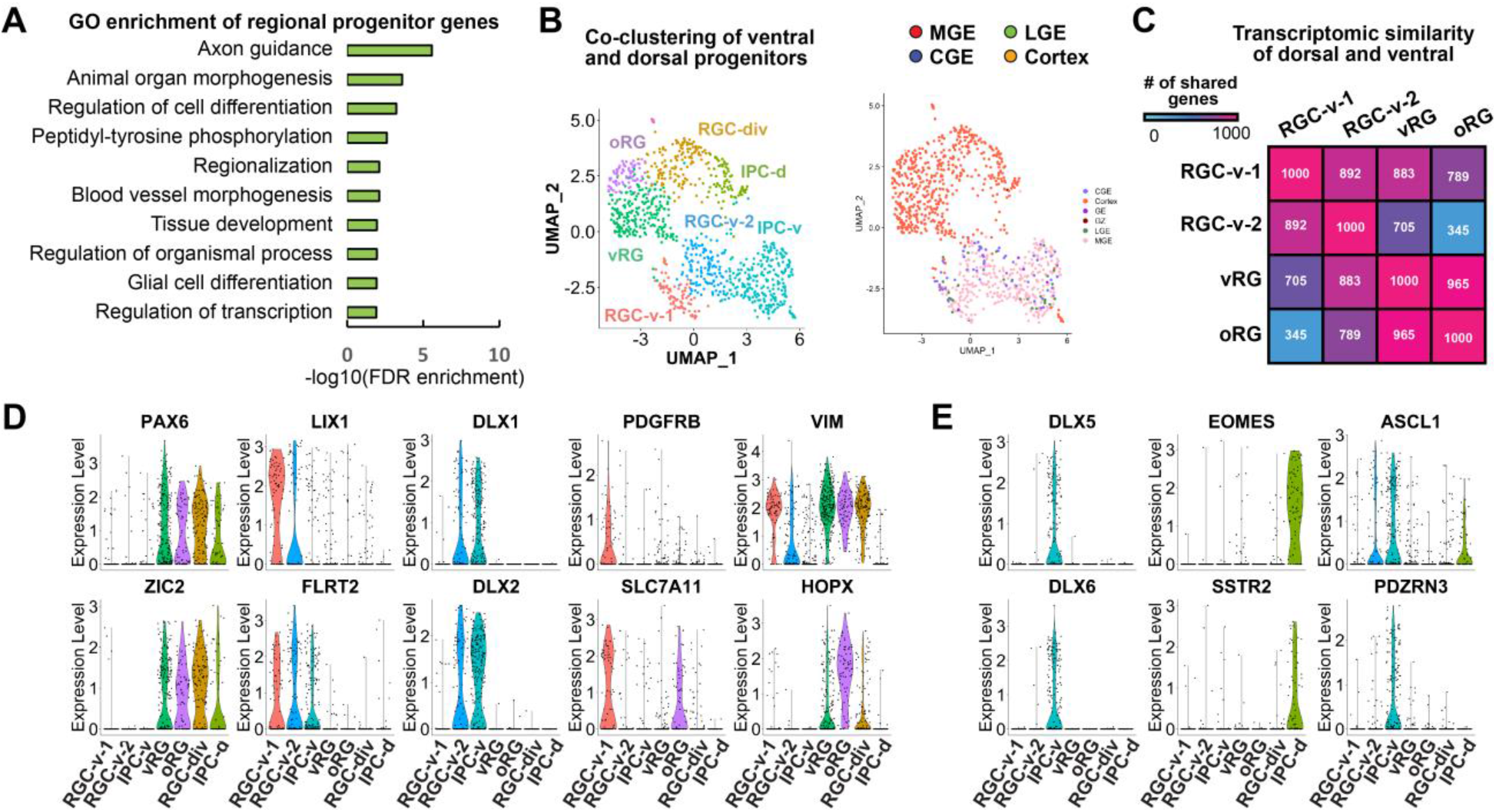
Comparative single-cell RNA-seq analysis of human and mouse cortical interneuron lineages. **A)** Gene ontology analysis of the genes enriched in the RGC and IPC populations of MGE and CGE. **B)** Coclustering of GE human progenitors with neural progenitor cells from developing human cortex. **C)** Transcriptomic similarity between the ventral and dorsal progenitor cell populations. **D)** Violin plots for genes enriched in specific subtypes of ventral and dorsal radial glia. **E)** Expression of ventral- and dorsal-specific IPC genes.

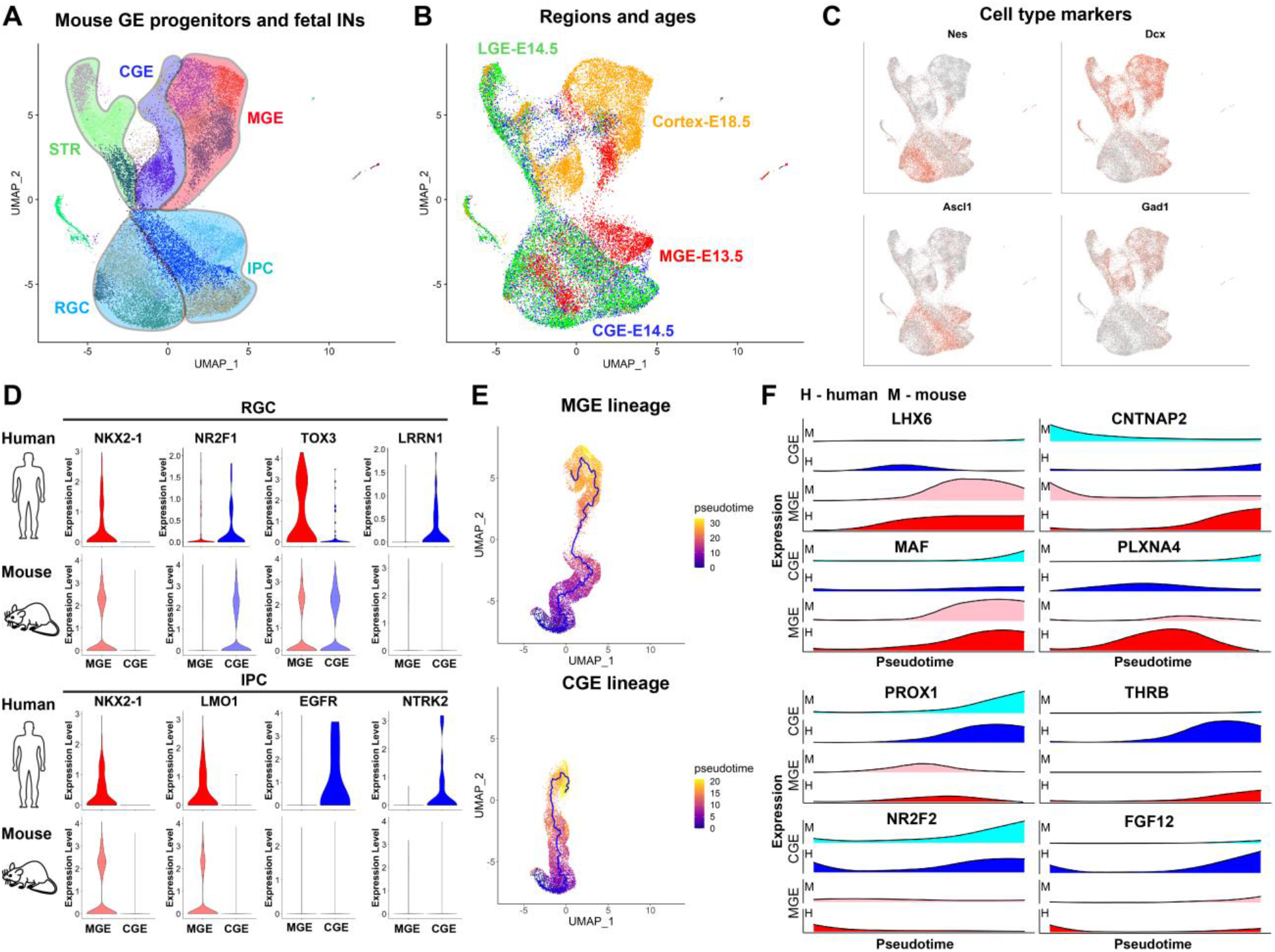
**A)** Co-clustering of mouse GE and cortical interneuron data during embryonic development. Clusters are grouped according to specific progenitor subtypes and lineages. **B)** Contribution of brain regions and developmental ages to transcriptomic clusters. **C)** Markers of GE progenitors and immature interneurons. **D)** Species-specific markers of radial glial and intermediate precursor cells expressed in the human or mouse GE. **E)** Comparison of MGE and CGE interneuron lineages between human and mouse. **F)** Lineagespecific genes shared and divergent between the two species.

**Figure S3.**
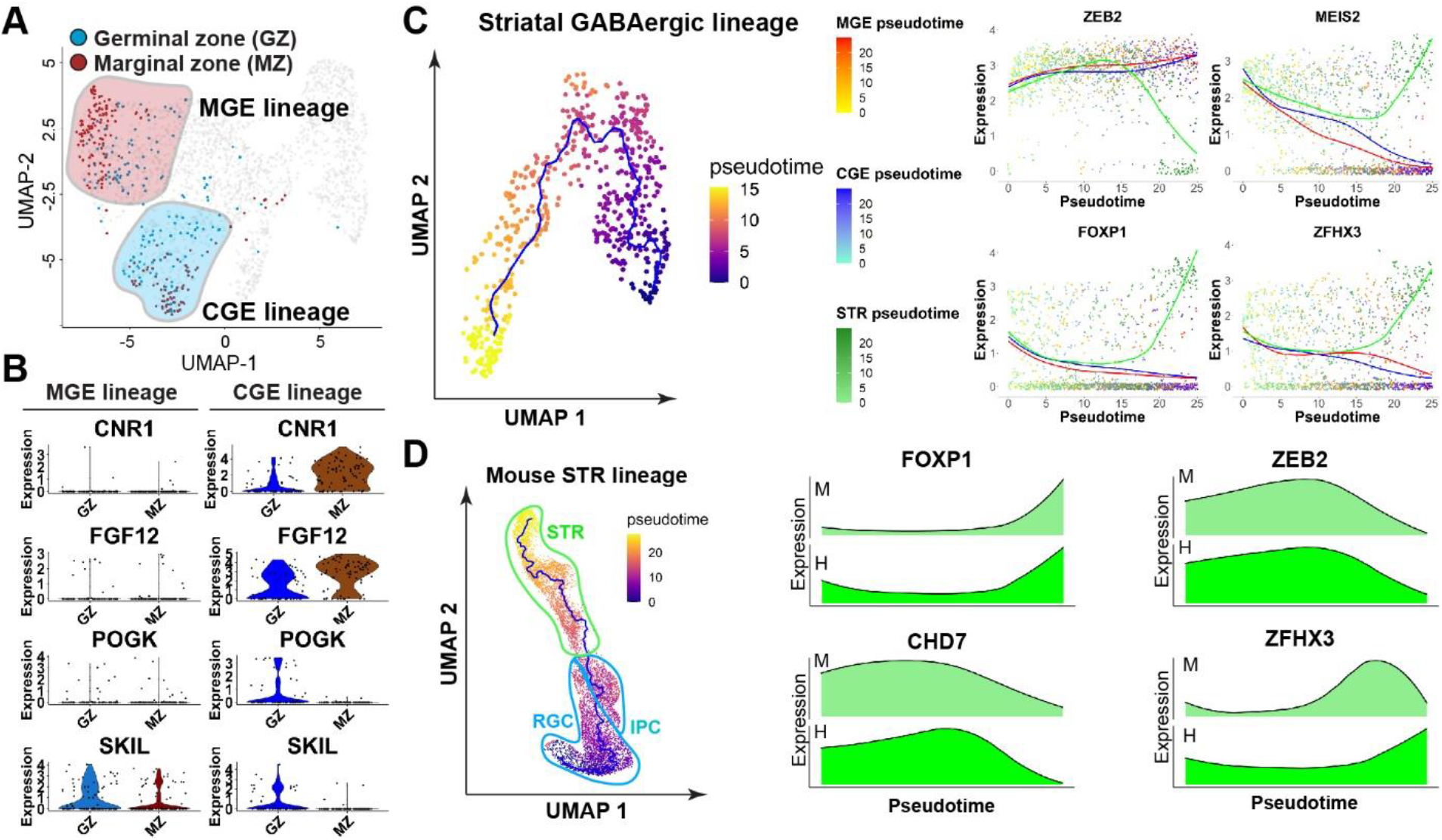
Analysis of the IN migratory streams and striatal GABAergic lineage. **A)** UMAP plot highlighting interneurons migrating through the cortical marginal (MZ) or germinal (GZ) zones. **B)** Genes specific to MZ or GZ migratory interneurons in the CGE lineage. **C)** Lineage trajectory and pseudotime for the human striatal GABAergic neuron lineage during the second trimester. Striatal-specific pseudotime-dependent genes are highlighted on the right. **D)** Trajectory analysis of the mouse STR lineage. STR-specific dynamically expressed genes that are common between the mouse and human are shown on the bottom.

**Figure S4.**
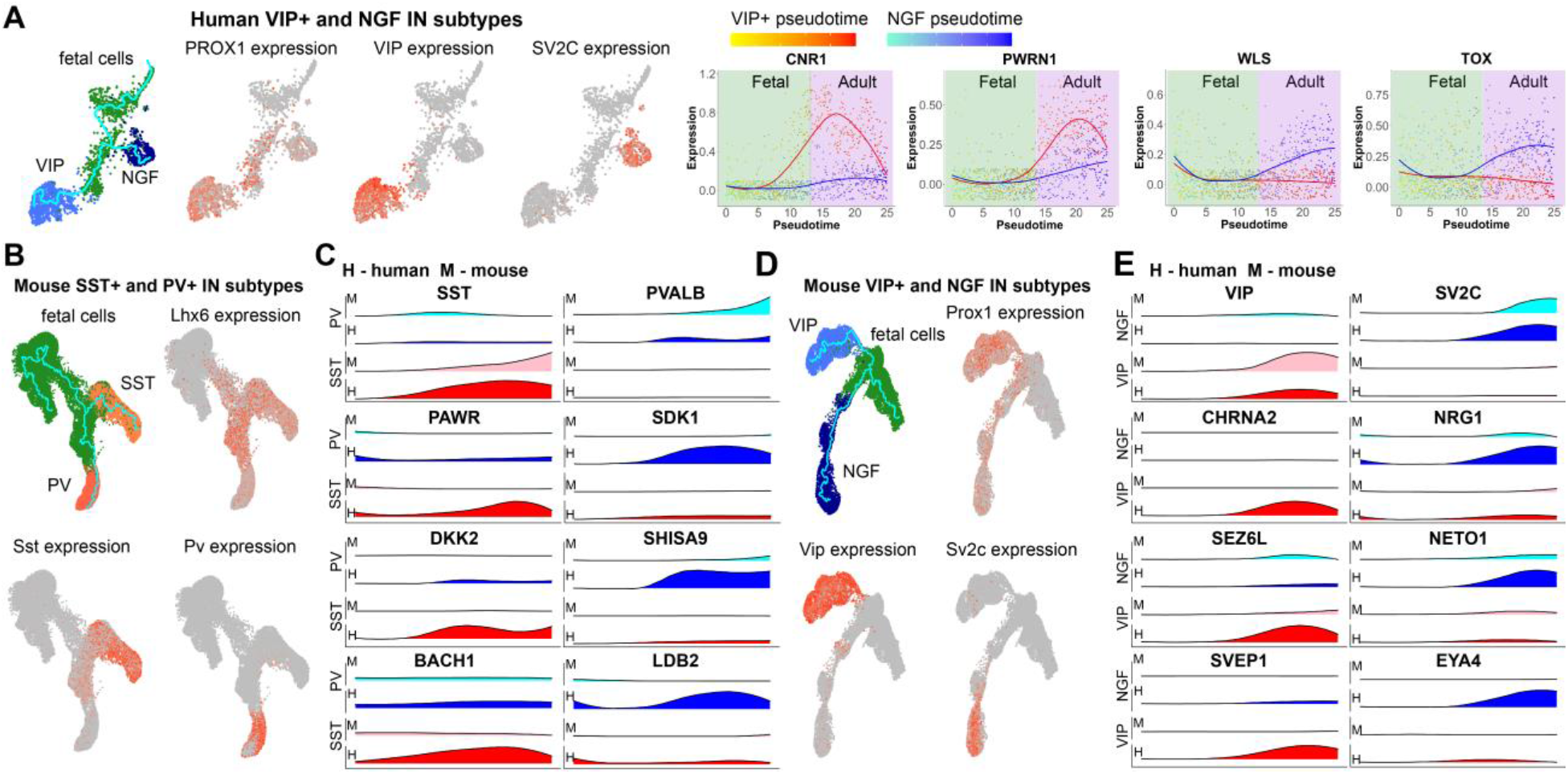
Comparison between human and mouse adult interneuron trajectories. **A)** Divergence of VIP and NGF CGE-derived interneuron populations as well as subtype-specific genes expressed during development. **B)** Trajectory analysis of mouse MGE-derived somatostatin (SST) and parvalbumin (PV) cortical interneurons. **C)** Canonical genes specifically expressed in human and mouse SST and PV interneurons (SST, PVALB, top row), as well as subtype-specific genes upregulated in human but not mouse lineages. **D)** Trajectory analysis of mouse CGE-derived VIP and NGF cortical interneurons. **E)** Genes specifically expressed in CGE-derived IN subtypes in both human and mouse cortex shown in top rows. Genes expressed in the human and not mouse are shown in lower rows.

**Table S1. Sample information and cell metadata.**

**Table S2. Markers of each transcriptomically defined cluster.**

**Table S3. Signatures of MGE and CGE RGC and IPC, as well as genes enriched in dorsal and ventral progenitors.**

**Table S4. Genes dynamically expressed during MGE, CGE and STR lineage development.**

**Table S5. Genes dynamically expressed during commitment to adult IN subtypes.**

**Table S6. List of antibodies used.**

## Methods

### Tissue samples

De-identified tissue samples were collected with previous patient consent in strict observance of the legal and institutional ethical regulations. Protocols were approved by the Human Gamete, Embryo, and Stem Cell Research Committee (institutional review board) at the University of California, San Francisco.

### Tissue processing

Tissue samples were dissected in artificial cerebrospinal fluid containing 125 mM NaCl, 2.5 mM KCl, 1mM MgCl_2_, 1 mM CaCl_2_, and 1.25 mM NaH_2_PO_4_under a stereotaxic dissection microscope (Leica). Samples for histology were fixed in 4% paraformaldehyde (PFA) prepared in calcium and magnesium free phosphate buffered saline (PBS) (pH^~^10) overnight at 4°C with constant agitation. PFA was then replaced with fresh PBS (pH=7.4) and cryopreserved by 24-48 hour incubation at 4oC in 30% sucrose diluted in PBS (pH=7.4) and embedded in a mixture of OCT (Tissue-Tek, VWR) and 30% sucrose. For single cell dissociation, tissue samples were further cut into small pieces and placed in a vial containing a pre-warmed solution of Papain (Worthington Biochemical Corporation) solution that was prepared according to the manufacturer protocol for 10min at 37°C. After approximately 60 min incubation, tissue was triturated following manufacturer protocol. After removing the dissociation media, cells were resuspended in PBS supplemented with 3% fetal bovine serum (Sigma) and quantified manually under a phase-contrast microscope. Samples were diluted to approximately 170,000 cells per ml before processing for scRNA-seq.

### Single cell RNA Sequencing

Single cell capture, cDNA synthesis and preamplification was performed as described before (Nowakowski et al., 2017b) using Fluidigm C1 auto-prep system following manufacturer’s protocol. Cells were stained using a live-dead kit (Life technologies) prior to capture of single cells, and each cell was imaged using a fluorescent microscope after the capture. Dead cells and multiplets were omitted from the downstream analysis. Library preparation was performed using the Illumina Nextera XT library preparation kit. Library concentration quantification was performed using Bioanalyzer (Agilent). Paired-end 100 bp sequencing was performed on the Illumina HiSeq2500. Raw scRNA-seq data can be found in dbGaP, accession number phs000989.

### Alignment

TrimGalore! 3.7 (https://github.com/FelixKrueger/TrimGalore) was used to trim 20 bp of adaptor sequencing, and paired-end alignments were performed using HISAT2 (Kim et al., 2015) to the human reference genome GRCh38. Counts for each cell were performed using subread-1.5.0 function featureCounts (Liao et al., 2014). After counts were obtained, normalization to counts per million was performed. Quality control was performed to further ensure that only high quality single cell data was processed further, and cells with fewer than 1000 genes/cell were removed. Only genes expressed in at least 5 cells were carried forward in the analysis.

### Single cell RNA-seq datasets

The following datasets were analyzed in this paper:

1. Human developing interneurons dataset, which included all cells captured from the ganglionic eminences (GE), as well as all MGE and LGE cells and all cells annotated as interneurons (INs) from Nowakowski et al (Nowakowski et al., 2017b). In addition, this dataset included cortical INs captured from the cortical germinal (GZ) and marginal (MZ) zones. The latter were defined as cortical interneurons by performing separate clustering with Seurat, annotating clusters based on IN marker expression and selecting cells in IN clusters.
2. Human neuronal progenitor dataset. This dataset included all cells from datasets (1) annotated as GE progenitors (radial glia cells, RGC, and intermediate precursor cells, IPC), as well as cortical radial glia and IPC from Nowakowski et al.
3. Mouse developing interneurons dataset. Raw count data for this dataset were obtained from Mayer et al (Mayer et al., 2018) and processed using the same pipeline for clustering and UMAP as the human data.
4. Human developing+adult interneurons dataset. This dataset was generated by combining dataset (1) and single-nuclei RNA-seq profiles generated on the 10x Genomics v.2 platform from neurotypical children and young adult individuals and annotated as cortical interneurons (Velmeshev et al., 2019). Single-cell and single-nuclei RNA-seq profiles were integrated using Seurat as detailed below.
5. Mouse developing+adult interneurons dataset. Raw count data from SMART-seq analysis of adult mouse cortex (Yao et al., 2020) was obtained from https://portal.brain-map.org/atlases-and-data/rnaseq/mouse-whole-cortex-and-hippocampus-smart-seq. Only data from cortical interneurons were analyzed. These data were integrated with dataset (3) using the Seurat integration approach.

### Integration of mouse single-cell RNA-seq and human single-nucleus RNA-seq datasets

To combine embryonic GE with cortex mouse scRNA-seq data in dataset (3), human prenatal scRNA-seq with postnatal snRNA-seq data in dataset (4) and developing and adult mouse scRNA-seq data in dataset (5), Seurat standard integration workflow and FindIntegrationAnchors and IntegrateData functions were used to perform data integration.

### Uniform Manifold Approximation and Projection (UMAP), clustering and cell type annotation

Version 3 of Seurat (Stuart et al., 2018) was used to perform principle component analysis, UMAP dimensionality reduction and Louvain clustering. In order to select the number of principle components (PCs) for UMAP and clustering, the scree plot method was used to choose the number of PCs after which the percentage of variance explained plateaus. Clustering was performed with the resolution parameter set to 1. In order to annotate cell types, feature plots were generated for known cell type and lineage markers.

### Identification of cell type and regional marker genes

To identify marker genes enriched in each cluster, `FindMarkers` function was used. Genes with adjusted P value < 0.05 and absolute log2(fold change) >= 0.5 were considered as enriched. The same approach was used to identify genes enriched in MGE and CGE radial glia and IPCs by comparing gene expression within the same cell type between the two regions using `FindMarkers` function. For the latter, all genes with adjusted P value < 0.05 were reported.

### Trajectory and pseudotime analysis

Monocle version 3 (Cao et al., 2019) was used to construct single-cell trajectories and calculate pseudotime. UMAP coordinates from the Seurat object were imported into Monocle using a custom R script (https://github.com/DmitryVel/Monocle3extension), and the trajectory graph was constructed using `learn_graph` function with the option `use_partition=F`. Then, to calculate pseudotime, `order_cells` function was used with the `root_pr_nodes` assigned to the starting node in the graph located in the radial glia cell cluster.

### Analysis of individual cell lineages

In order to select individual branches of the graph associated with specific lineages, a custom R script (https://github.com/DmitryVel/Monocle3extension) was used to perform the following steps: 1) Select a non-branching segment of the graph between the starting node in the radial glia cell cluster and the terminal node in the mature IN cluster using `all_simple_paths` from igraph package. 2) In case multiple segments are selected between the two nodes, select the one passing through intermediate clusters expressing lineage-specific markers (e.g., for MGE lineage, select branches passing through clusters expressing LHX6). 3) Calculate average Euclidian distance between adjacent nodes of the selected branch (internode distance). 4) Select cells along the branch that are no farther than 5 times of the internode distance from one of the nodes in the branch.

### Identification of lineage-specific dynamically expressed genes

To identify pseudotime-correlated genes in each lineage, Monocle 3’s `graph_test` was used on cells selected in each lineage to perform Moran’s I test. Genes with adjusted P value < 0.05 and Moran’s I >=0.15 were considered as significantly correlated with pseudotime. To identify genes dynamically expressed in each lineage, we calculated the difference of Moran’s I between lineages (delta I). Genes with delta I >= 0.15 were prioritized as most lineage-specific genes. To further classify genes in individual lineages based on their dynamic expression pattern, we performed time series clustering for genes from both the human MGE and CGE lineages using TSclust R package (Montero and Vilar, 2014). We calculated distance using the temporal correlation and raw values (CORR) method and performed hierarchical clustering using ward.D2 method. To determine the optimal number of clusters, we used the silhouette method. To construct averaged curves for each cluster, we averaged expression of all genes in each temporal cluster at each pseudotime point.

### Pseudotime gene expression plots

To plot pseudotime-dependent expression of selected genes in two lineages simultaneously, we used the following approach: 1) Compress gene expression estimates for each gene along pseudotime using a sliding window of width = 3 and step size equal to 500/(number of pseudotime points). 2) Fit pseudotime expression curves using the GLM approach, 3^rd^ degree polynomial and quasipoisson distribution. Expression values were log10 transformed prior to fitting. 3) Smooth the resulting curves using the loess procedure and polynomial degree of 2. The resulting transformed expression and pseudotime values and fitted curves can then be plotted on the same scale. For plotting mouse and human lineages side-by-side, same procedure was used but a ridgeline plot was constructed instead.

### Gene ontology analysis

PANTHER (Thomas et al., 2003) was used to perform statistical overrepresentation test using the GO biological process complete classification. All genes expressed in the cell type or lineage subjected to GO analysis were used as the reference list. All GO processes with adjusted enrichment P value < 0.05 were considered significant. Only the processes on the top of PANTHER process hierarchy were reported.

### Immunohistochemistry analysis

Frozen slides were allowed to equilibrate at 4⍰C overnight and then to room temperature for 3 h. After baking at 60 °C for 30 min, slides were placed in 4% PFA for 15 min, rinsed with TNT buffer (0.05% Triton-X100 in PBS) and water, and then placed in 3% H2O2 in PBS for 30 min. Slides were rinsed in TNT buffer and water, and antigen retrieval was conducted for 10 min at 95 °C in 10 mM sodium citrate buffer, pH 6.0. Following antigen retrieval, slides were allowed to cool to 75 °C, rinsed with TNT buffer and water, and placed in 3% H2O2 in PBS for 90 min. After three 10 min TNT buffer washes, slides were blocked with TNB solution (0.1 M Tris-HCl, pH 7.5, 0.15 M NaCl, 0.5% blocking reagent from PerkinElmer) for 1 h. Slides were incubated in primary antibody overnight at 4 °C and in biotinylated secondary antibody (Jackson Immunoresearch Laboratories) for 2.5 h at room temperature. All antibodies were diluted in TNB solution. For most antibodies, the conditions of use were validated by the manufacturer (antibody product sheets). When this information was not provided, we performed control experiments, including no primary antibody (negative) controls and comparison to mouse staining patterns.

After three 10 min TNT buffer washes, sections were incubated for 30 min in streptavidin–horseradish peroxidase, which was diluted (1:200) with TNB. Slides were then rinsed in three 5 min TNT buffer washes. Tyramide signal amplification (PerkinElmer) was used for all antigens. Sections were incubated in tyramide-conjugated fluorophores for 10 min at the following dilutions: fluorescein: 1:100; Opal 570: 1:100; Opal 520, and for 5 min at the following dilution: fluorescein: 1:100; Cy5. For multichannel stains, sections were then rinsed several times in TNT buffer, placed in 4% PFA for 15 min, rinsed in TNT buffer and water, and then placed in 3% H2O2 in PBS for 2 h. After three 10 min TNT buffer washes, slides were blocked with TNB solution for 30 min to 1 h, and incubated with the next primary antibody at 4 °C overnight. After repeating the secondary antibody incubation, streptavidin-horseradish peroxidase incubation, and Tyramide signal amplification steps, sections were rinsed several times in TNT buffer, dehydrated, mounted and coverslipped after the last channel development.

List of antibodies used in available in Table S6.

### Single molecule fluorescent *in situ* hybridization combined with Immunohistochemistry

Single molecule fluorescent *in situ* hybridization (smFISH) and immunohistochemistry were performed on fixed frozen sections from human brain specimen of indicated ages using Advanced Cell Diagnostics (ACDBio) RNAscope^®^ Fluorescent Multiplex Reagent Kit and probes. Cryosections (14-16μm thick) were mounted on glass slides and washed in RNase free PBS for 5 mins. Slides were first baked at 60°C for 20 mins and the sections were dehydrated in 50%, 70% and 100% ethanol for 5 mins each at room temperature and then air dried. Using RNAscope reagents, target retrieval was performed for 5 mins at 95 °C and sections were washed with distilled water. Slides were then placed in 1% H_2_O_2_ in PBS for 45 minutes and were blocked with Tris-NaCl blocking buffer (0.1 M Tris-HCl, 0.15 M NaCl and 0.5%(w/v) blocking reagent from PerkinElmer). Slides were incubated in primary antibodies overnight at 4°C followed by three washes in PBS-T for 2 mins each at room temperature. A post-primary fixation step was performed in 4% paraformaldehyde at 4°C, followed by an additional wash with PBS-T for 2 mins. Sections were next treated with Protease IV reagent for 30 mins at 40°C and were maintained in RNase free water until hybridization step. RNAscope probes were diluted at 1:50 ratio in channel 1 probe and preheated to 40°C for 5 mins. Slides were incubated with this probe mix for 2 hours at 40°C. After probe hybridization, slides were washed twice for 2 mins each (in 1x RNAscope Wash Buffer Reagent from ACDBio) before proceeding to the fluorescent detection step according to manufactures protocol. Briefly, sections were incubated in AMP 1-FL for 30 min at 40°C, washed twice, incubated in RNAscope AMP 2-FL for 15 min at 40°C, washed twice, incubated in RNAscope AMP 3-FL for 30 min at 40°C, washed twice and incubated in fluorophore containing AMP 4-FL solution for 15 mins at 40°C and washed twice. Slides were next incubated in biotinylated secondary antibody for 1 hour at room temperature followed by incubation in fluorophore tagged horseradish peroxidase (Alexa Fluor; Thermofisher). Sections were washed thrice in PBS-T and counterstained with RNAscope DAPI for 30 seconds. Slides were mounted using prolong gold antifade. Images were acquired at 63x on Lecia SP8 upright AOBS confocal microscope.

