## Supplementary information for "Molecular diversity and lineage commitment of human interneuron progenitors"

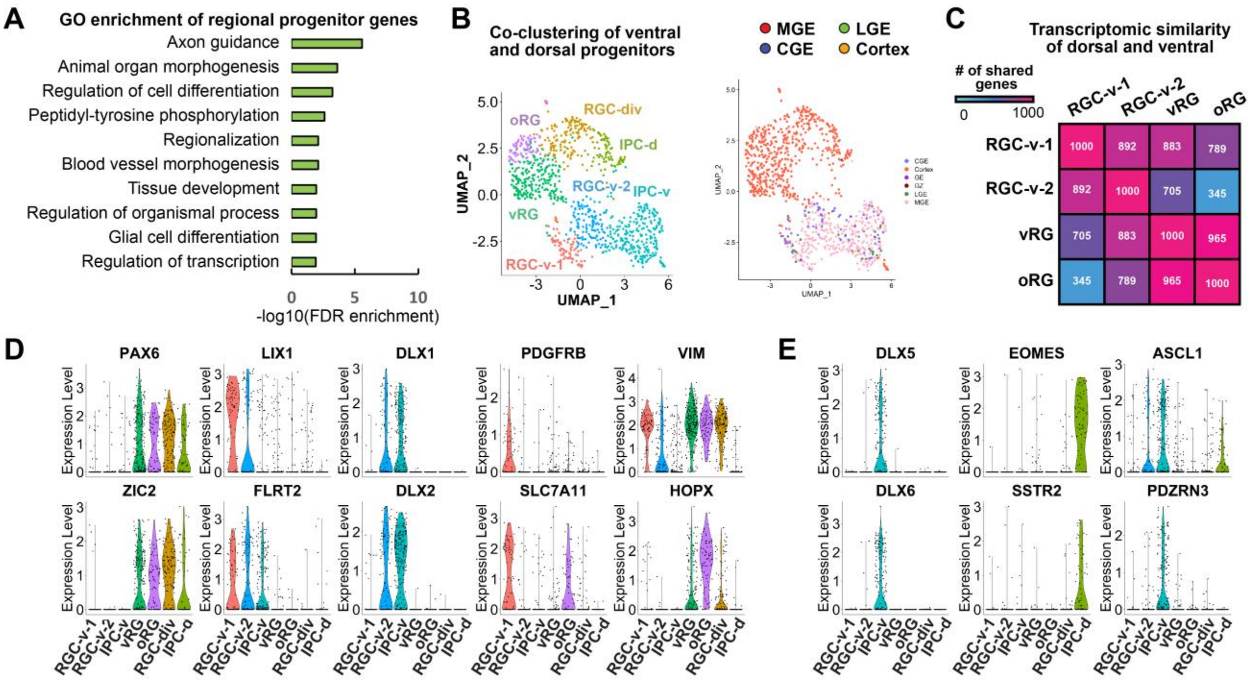


**Figure S1. Analysis of human ventral and dorsal neural progenitor subtypes.**

**A)** Gene ontology analysis of the genes enriched in the RGC and IPC populations of MGE and CGE. **B)** Co-clustering of GE human progenitors with neural progenitor cells from developing human cortex. **C)** Transcriptomic similarity between the ventral and dorsal progenitor cell populations. **D)** Violin plots for genes enriched in specific subtypes of ventral and dorsal radial glia. **E)** Expression of ventral- and dorsal-specific IPC genes.

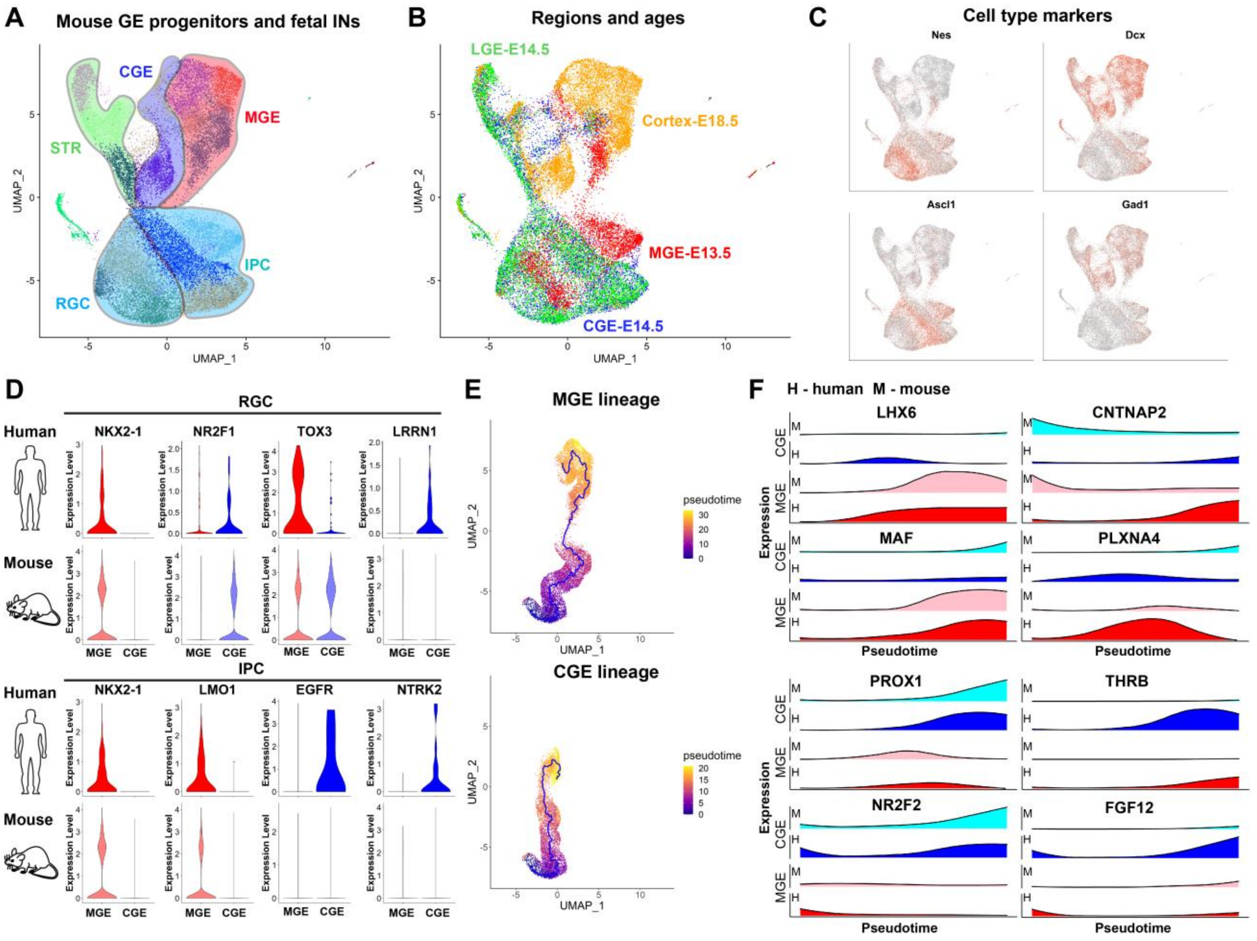


**Figure S2. Comparative single-cell RNA-seq analysis of human and mouse cortical interneuron lineages.**

**A)** Co-clustering of mouse GE and cortical interneuron data during embryonic development. Clusters are grouped according to specific progenitor subtypes and lineages. **B)** Contribution of brain regions and developmental ages to transcriptomic clusters. **C)** Markers of GE progenitors and immature interneurons. **D)** Species-specific markers of radial glial and intermediate precursor cells expressed in the human or mouse GE. **E)** Comparison of MGE and CGE interneuron lineages between human and mouse. **F)** Lineage-specific genes shared and divergent between the two species.

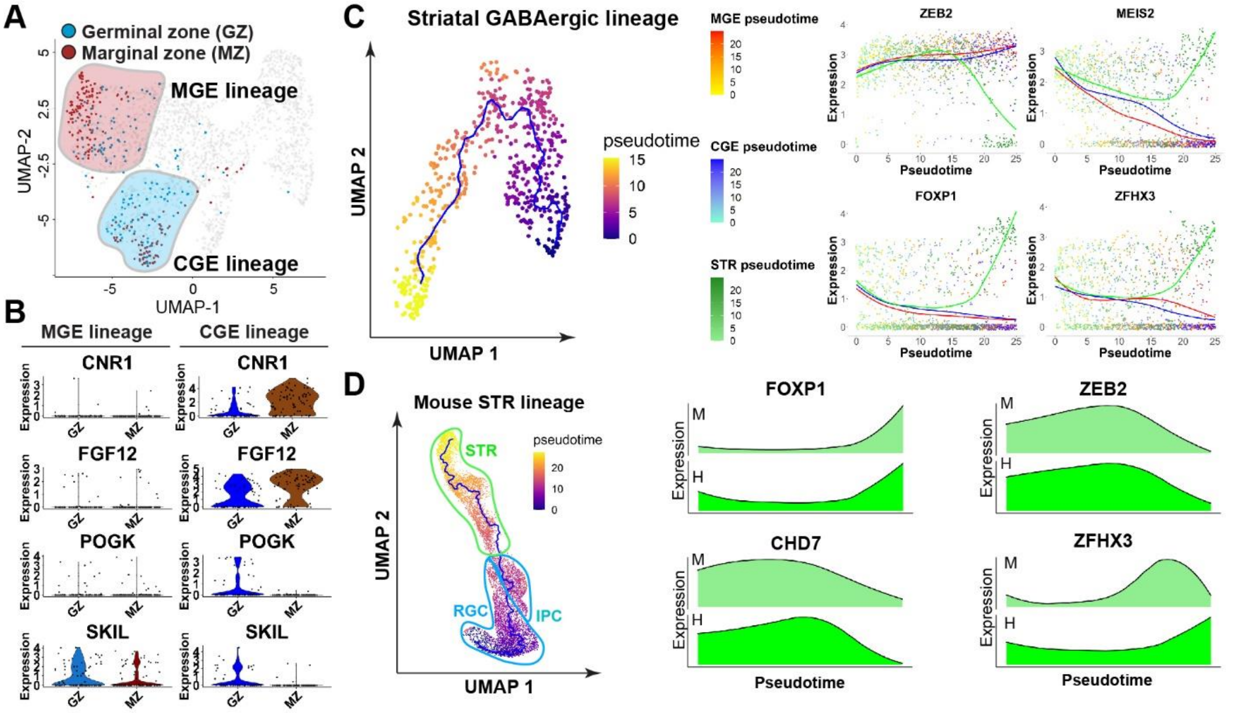


**Figure S3. Analysis of the IN migratory streams and striatal GABAergic lineage. A)** UMAP plot highlighting interneurons migrating through the cortical marginal (MZ) or germinal (GZ) zones. **B)** Genes specific to MZ or GZ migratory interneurons in the CGE lineage. **C)** Lineage trajectory and pseudotime for the human striatal GABAergic neuron lineage during the second trimester. Striatal-specific pseudotime-dependent genes are highlighted on the right. **D)** Trajectory analysis of the mouse STR lineage. STR-specific dynamically expressed genes that are common between the mouse and human are shown on the bottom.

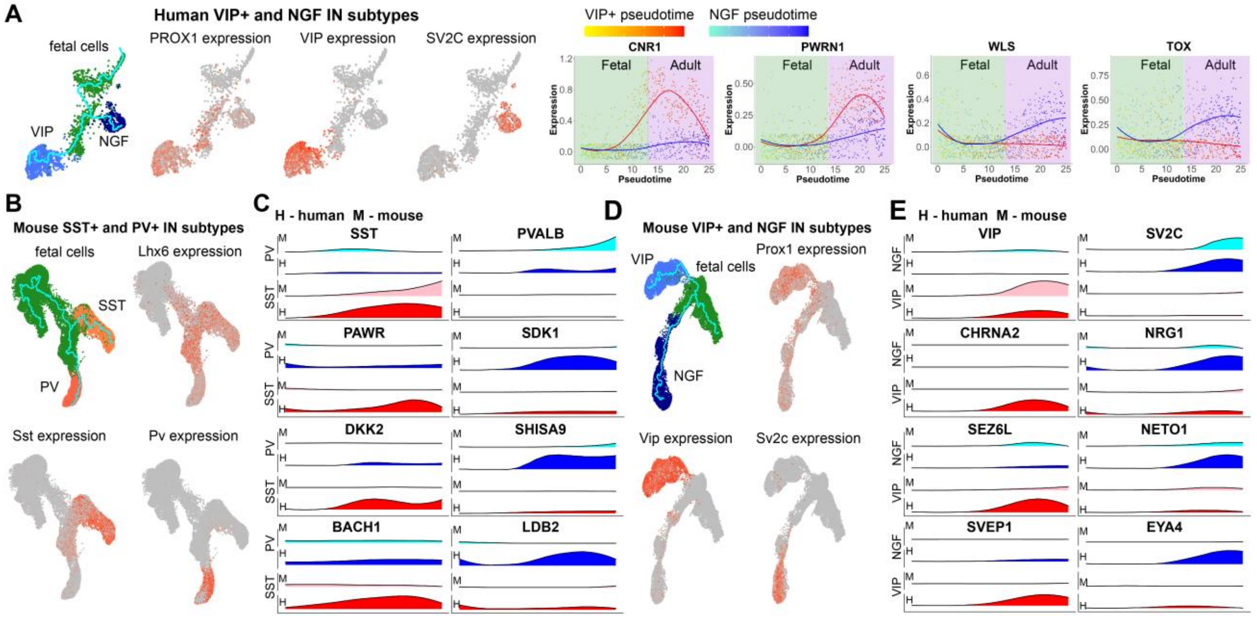

**Figure S4. Comparison between human and mouse adult interneuron trajectories. A)** Divergence of VIP and NGF CGE-derived interneuron populations as well as subtype-specific genes expressed during development. **B)** Trajectory analysis of mouse MGE-derived somatostatin (SST) and parvalbumin (PV) cortical interneurons. **C)** Canonical genes specifically expressed in human and mouse SST and PV interneurons (SST, PVALB, top row), as well as subtype-specific genes upregulated in human but not mouse lineages. **D)** Trajectory analysis of mouse CGE-derived VIP and NGF cortical interneurons. **E)** Genes specifically expressed in CGE-derived IN subtypes in both human and mouse cortex shown in top rows. Genes expressed in the human and not mouse are shown in lower rows.
